# Low level of expression of C-terminally truncated human FUS causes extensive changes in spinal cord transcriptome of asymptomatic transgenic mice

**DOI:** 10.1101/689414

**Authors:** Ekaterina A. Lysikova, Sergei Funikov, Alexander P. Rezvykh, Kirill D. Chaprov, Michail S. Kukharsky, Alexey A. Ustyugov, Alexey V. Deykin, Ilya. A. Flyamer, Shelagh Boyle, Sergey O. Bachurin, Natalia Ninkina, Vladimir L. Buchman

## Abstract

Mutations in a gene encoding RNA-binding protein FUS was linked to familial forms of amyotrophic lateral sclerosis (ALS). C-terminal truncations of FUS are associated with aggressive forms of ALS. However, motor neurons are able to tolerate permanent production of pathogenic truncated form of FUS protein until its accumulation in the cytoplasm of neurones does not reach a critical threshold.

In order to identify how the nervous system responds to pathogenic variants of FUS we produced and characterised a mouse line, L-FUS[1-359], with a low level of neuronal expression of a highly aggregation prone and pathogenic form of C-terminally truncated FUS. In contrast to mice with substantially higher level of expression of the same FUS variant that develop severe early onset motor neuron pathology, L-FUS[1-359] mice do not develop any sign of pathology even at old age. Nevertheless, we detected substantial changes in the spinal cord transcriptome of these mice comparing to the wild type littermates. We suggest that at least some of these changes reflect activation of cellular mechanisms compensating to potentially damaging effect of pathogenic FUS production. Further studies of these mechanism might reveal effective target for therapy of FUS-ALS and possibly, other forms of ALS.

## Introduction

Mutations in the gene encoding FUS are among the most frequent causes of familial ALS (fALS) and many of these mutations cause partial mislocalisation of this protein from the nucleus to the cytoplasm of motor neurons where it aggregates with formation of large inclusions and compromise various important cell functions ^1-4^. Moreover, aggregation of FUS has been observed in the cytoplasm of neurons in a subset of patients with sporadic form of ALS (sALS) as well as in patients with certain variants of frontotemporal lobar degeneration (FTLD-FUS) in the absence of any mutations or polymorphisms of the gene encoding FUS ^5-9^.

Various invertebrate and vertebrate animal models of FUS malfunction, including several transgenic FUS mouse lines have been produced and characterised in order to shed light on molecular mechanisms of ALS-FUS and FTLD-FUS but the results of these studies were not always consistent (reviewed in ^10-12^). This might reflect a complex combination of intrinsic (e.g. even subtle differences in genetic background) and environmental factors (e.g. type of animal diet) that affect the manifestation of pathology in animals with genetic alterations of FUS structure of expression. However, these are obvious similarities in how pathological changes develop in all animal models. Motor neurons of transgenic mice expressing pathogenic variants of human FUS as well as motor neurons of ALS patients carrying germ line mutations of FUS survive and function normally for a relatively long period of time, suggesting that they successfully employ certain intracellular mechanisms preventing damaging effects of the pathogenic protein when it present at low levels. An ability to artificially boost these mechanisms activity, either directly or via modulation of their regulatory pathways, can help counteracting pathogenic effects of FUS and potentially, other ALS-linked proteins even when they accumulate in the neuronal cytoplasm beyond a threshold that overload an intrinsic ability of the neuronal defence systems. Therefore, it is important to identify how the nervous system responds to low levels of pathogenic variants of FUS and prevents potentially damaging consequences of its permanent production.

Previously we have described production and characterisation of a transgenic mouse line FUS 1-359 clone 19 (S-FUS[1-359]) with neurospecific expression of C-terminally truncated form of human FUS (tr-hFUS) lacking its nuclear localisation signal and RGG domains ^13^. This modification recapitulates consequences of FUS gene mutations in certain forms of familial ALS ^14-17^ typified by mislocalisation of modified FUS and aggressive disease with young onset and rapid progression ^18^. Although heterozygous mice of S-FUS[1-359] line express tr-hFUS at the lower level than the level of expression of the endogenous mouse FUS, accumulation of tr-hFUS in the cytoplasm of neurons leads to its aggregation and the development of FUS-proteinopathy characterised by the presence of large inclusions, neuronal dysfunction and death. These pathological changes are most pronounced in the motor neurons of the spinal cord, which causes the development of severe and fast progressing motor neuron disease in young adult S-FUS[1-359] mice and their death within one to three weeks from the onset of clinical signs ^13^.

Here we assessed changes in the spinal cord transcriptome of L-FUS[1-359] mice, another transgenic mouse line produced along to S-FUS[1-359] line using the same transgenic construct. Although these mice have substantially lower level of tr-hFUS expression in the spinal cord and do not develop clinical signs of motor neuron pathology, we found a significant number of transcripts that differentially regulated in the spinal cord of L-FUS[1-359] mice when compared to their wild type littermate.

## Results

### Transgenic mice expressing low level of C-terminally truncated human FUS in their neurons do not develop motor neuron pathology

A founder of L-FUS[1-359] transgenic mouse line was produced by pronucleus microinjection of the same transgenic construct as was used for production of the previously described S-FUS[1-359] line, namely a linear DNA fragment containing human FUS cDNA encoding amino acids 1 to 359 (tr-hFUS) under control of regulatory elements of mouse Thy-1 gene ^13^. The L-FUS[1-359] line was established on a CD1 genetic background by serial backcrosses (>5) with the wild type CD1 mice. Intercrosses of heterozygous animals produced homozygous L-FUS[1-359] mice and their wild type littermates that were used in some of experiments described below. Analysis of genomic DNA revealed that L-FUS[1-359] mice carry the same number of tandemly arranged repeats of the transgenic cassette as S-FUS[1-359] mice (Suppl. Fig. S1). However, the localisation on different chromosomes (Suppl. Fig. S1) and presumably, different chromatin organisation around the integration sites led to substantially different expression levels of the transgenic cassette in the neural tissues of these two mouse lines. Quantitative RT-PCR revealed that the level of tr-hFUS mRNA in the spinal cord of 9-week old L-FUS[1-359] mice is 20 times lower than in the spinal cord of presymptomatic S-FUS[1-359] of the same age and maintained at the same CD1 genetic background (Fig. 1a). Consistently, Western blot analysis with human FUS specific antibody revealed substantially lower level of tr-hFUS protein in the spinal cords of L-FUS[1-359] mice when compared to S-FUS[1-359] mice (Fig. 1b). While accumulation of tr-hFUS in the cytoplasm of spinal motor neurons of S-FUS[1-359] mice leads to its aggregation with formation of large cytoplasmic inclusions at the onset of clinical signs, i.e. at the age of 13 to 20 weeks (Fig. 1c and 13), tr-hFUS remains diffusely distributed in the cytoplasm and also present in the nucleus of spinal motor neurons of L-FUS[1-359] mice (Fig. 1c). The latter observation is consistent with only partial cytoplasmic mislocalisation of FUS carrying point mutations in or completely lacking the NLS reported in neurons of mice expressing low levels of these human FUS variants ^38-40^.

**Figure 1.**
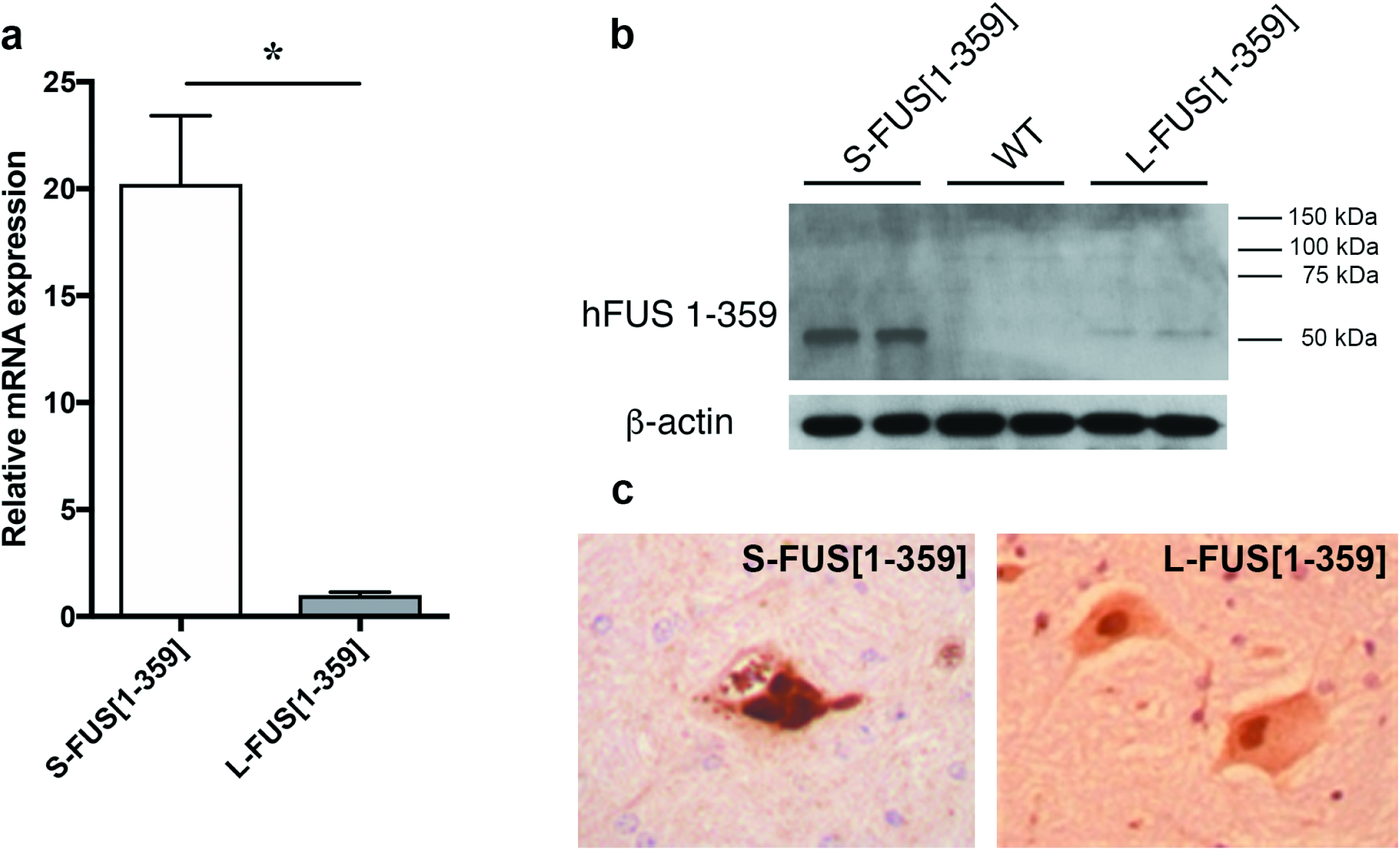
Expression of tr-hFUS in the thoracic spinal cord of L-FUS[1-359] and S-FUS[1-359] mice. a) Relative mRNA levels quantified by real-time qRT-PCR with primers specific for human FUS mRNA. Total RNA samples from four 9-week old hemizygous male animals were analysed in triplicates (*p<0.01, Mann-Whitney U-test). b) Western blot analysis of total protein samples extracted from spinal cords of two hemizygous male L-FUS[1-359], two hemizygous male S-FUS[1-359] and two wild type mice (all 9-week old) with antibody specific for human FUS. For loading control the same membrane was re-probed with anti-beta-actin antibody. Full-length blots provided in Suppl. Fig. S3. c) Immunohistochemical staining of transverse section through the spinal cord of 18-month old hemizygous male L-FUS[1-359] mouse and 4-month old early symptomatic hemizygous male S-FUS[1-359] with antibody specific for human FUS. Note accumulation of tr-hFUS in large cytoplasmic and small nuclear inclusions in an anterior horn motor neuron of the S-FUS[1-359] mouse and diffuse distribution of tr-hFUS in the cytoplasm and the nucleus of the L-FUS[1-359] mouse. Scale bars, 50 µm.

The low level of tr-hFUS expression is not sufficient for triggering motor neuron pathology or any other obvious disease phenotypes even in homozygous L-FUS[1-359] mice (Fig. 2). The lifespan of these animals was not different from the lifespan of their littermates (Suppl. Fig. S2) and even in ageing animals no sign of spinal motor neuron loss was noted (Fig. 3).

**Figure 2.**
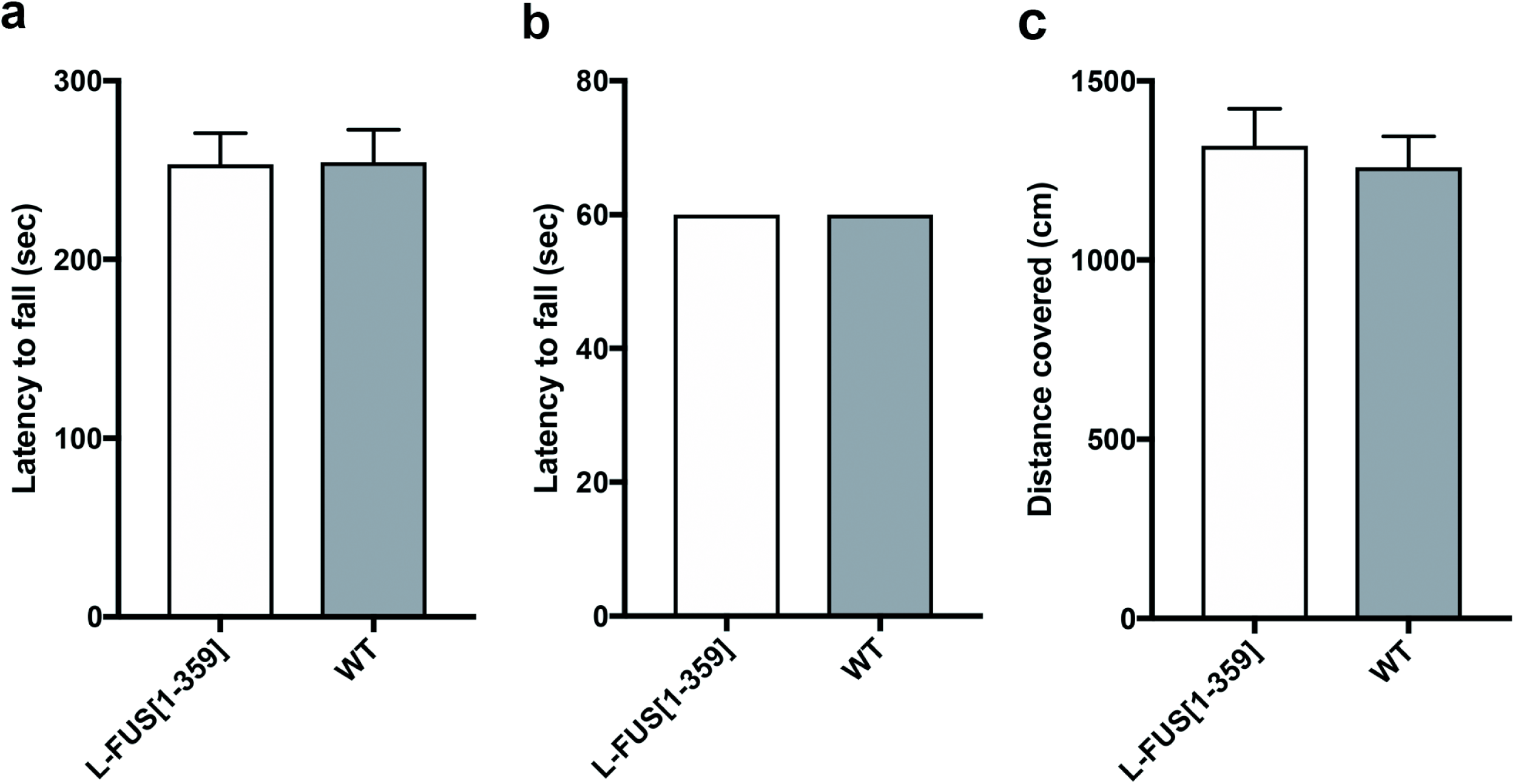
Animal performance in motor behaviour tests and their locomotor activity in novel non-anxiogenic environment. Bar charts show means±SEM of values obtained by testing one-year old homozygous male L-FUS[1-359] and wild type (WT) mice (10-12 per genotype). a) The latency to fall from the accelerating rotarod. No significant difference between performance of two groups of mice were found (p>0.05, Mann-Whitney U-test). b) The latency to fall from the inverted grid. Note that all animals in each group successfully completed this 60-second test. c) Total distance covered by an animal over the 5-minute test in the activity camera. No significant difference between performance of two groups of mice were found (p>0.05, Mann-Whitney U-test).

**Figure 3.**
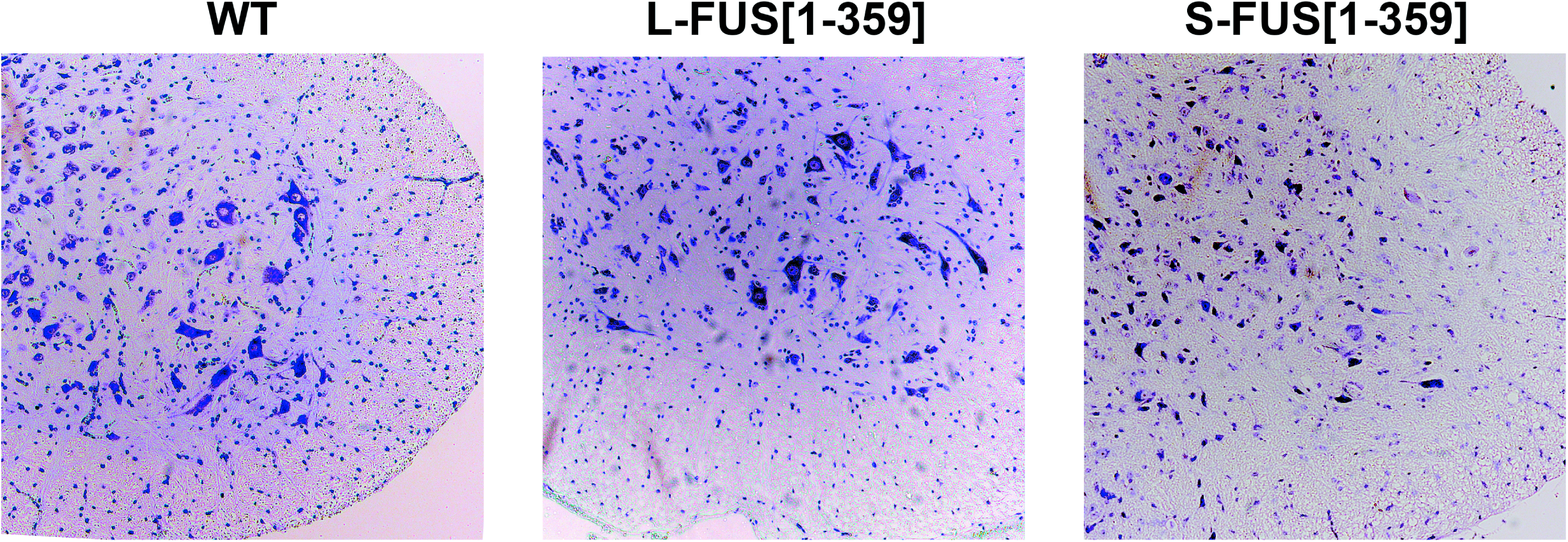
No loss of spinal motor neurons in ageing L-FUS[1-359] mice. Nissl stained transverse section through the spinal cord of 18-month old hemizygous male L-FUS[1-359] mouse, its wild type littermate and 5-month old symptomatic hemizygous male S-FUS[1-359] mouse. Note that while few motor neurons remained in the anterior horn of S-FUS[1-359] mice are either pale stained or shrinked, the anterior horn of ageing L-FUS[1-359] mice have the same complement and morphology of motor neurons as anterior horn of wild type animals.

### Changes of spinal cord transcriptomes caused by expression of C-terminally truncated human FUS

To assess whether a low level of expression of a pathogenic variant of FUS affects gene expression in the spinal cord of transgenic mice we employed RNA sequencing to compare spinal cord transcriptomes of four 9-week old hemizygous L-FUS[1-359] mice and four of their wild type littermates. As a result of deep-sequencing, we obtained from 13 to 17 million reads for each library produced from an individual mouse thoracic spinal cord.

Our analysis of sequencing data revealed 272 DEGs with FDR < 0.05 that clearly distinguish gene expression profiles of L-FUS[1-359] and WT mice on multidimensional scaling (Fig. 4a, b, Supplementary table S1). In order to discriminate DEGs that are intrinsic for neurons and glial cells we used the published datasets of purified microglia and laser-microdissected ventral horns of mice spinal cord applying a 5-fold change cutoff of gene expression levels ^30,31^. As a result, we obtained 200 genes (> 73% of DEG, FDR < 0.05) that are more specific for spinal cord neurons than for microglial cells and only 7 genes (< 3% of DEG, FDR < 0.05) that are intrinsic to microglia (Fig. 4c). Remaining 65 genes (∼ 24% of DEG) were shared by both types of cells (Fig. 4c). Thus, the observed gene expression changes in the spinal cord of L-FUS[1-359] mice take place predominantly in neurons.

**Figure 4.**
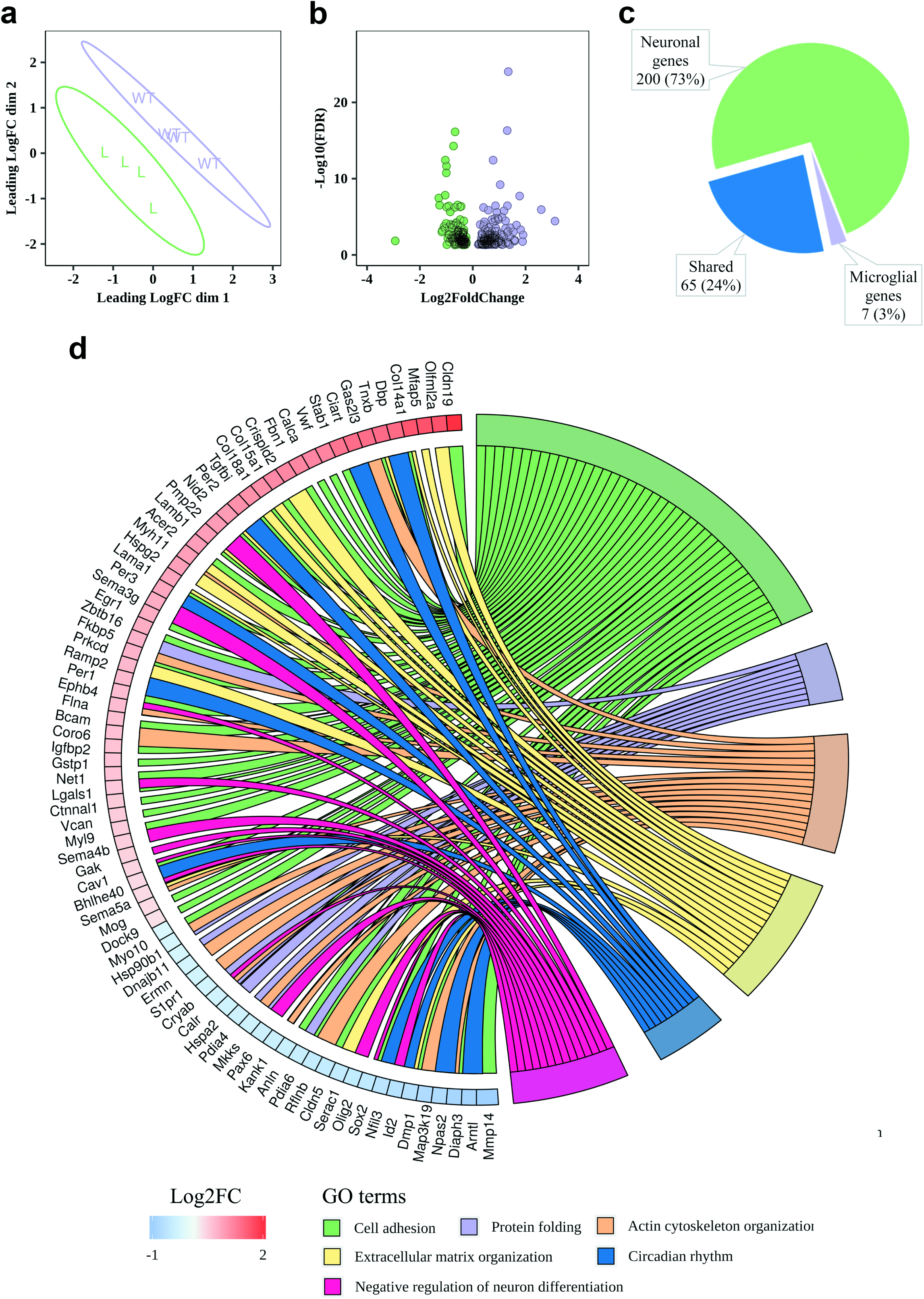
Gene expression profile changes in the spinal cord of L-FUS(1-359) mice compared to wild type littermates. a) Multidimensional scaling plot of RNA sequencing data for L-FUS[1-359] mice (L, in purple) and wild type mice (WT, in green). b) Volcano plot demonstrating an amount of DEGs in the spinal cord of L-FUS[1-359] mice compared to wild type mice (FDR < 0.05). c) Distribution of DEGs between neuronal and microglial cells in the studied samples. The sorting of genes to microglia and neurons was performed using a 5-fold change cutoff of gene expression levels (FDR < 0.05). d) Gene ontology analysis of DEGs. The most enriched GO terms of biological processes are present (p < 0.05) and sorted by their expression level (Log2FC, FDR < 0.05) in L-FUS[1-359] mice in comparison to WT mice.

The gene ontology (GO) enrichment analysis of genes differentially expressed in L-FUS[1-359] spinal cord revealed several groups of genes encoding proteins involved in biological processes important for normal function of the nervous system. The largest group includes upregulated genes for claudin 19 (*Cldn19*), tenascin XB (*Tnxb*), stabilin 1 (*Stab1*), basal cell adhesion molecule (*Bcam*), collagens (*Col14a1, Col15a1, Col18a1*), collagen-binding proteins (*Tgfbi, Nid2*), semaphorins (*SEMA3G, SEMA4B* and *SEMA5A*) and several other genes involved in cell adhesion and organisation of extracellular matrix (green and yellow groups in Fig. 4d).

A significant number of genes involved in negative regulation of neuronal differentiation is also upregulated in the spinal cord of L-FUS[1-359] mice, however genes encoding transcriptional factor SOX2 and OLIG2 that play a central role in neuronal cell differentiation and spinal motor neuron specification, respectively are significantly downregulated (red group in Fig. 4d).

The circadian rhythm related genes are highly represented between DEGs – all three paralogs of the “core” clock activator gene *period* – *Per1, Per2* and *Per3*, genes encoding the circadian transcriptional repressors CIART and BHLHE40, and transcription factor DBP are upregulated in the spinal cord neurons of L-FUS[1-359] mice, whereas genes encoding transcriptional activators ARNTL and NPAS2, and transcriptional repressor NFIL3 are significantly downregulated (blue group in Fig. 4d).

Expression of genes encoding the proteins assisting protein folding in the endoplasmic reticulum such as chaperones and chaperonins *Hspa2, Hsp90b1, Dnajb11, Cryab, Mkks*, protein disulfide isomerases *Pdia4, Pdia6* as well as calreticulin (*Calr*) are decreased in the spinal cord of L-FUS[1-359] mice (lilac group in Fig. 4d).

We also observed the altered expression levels of genes encoding several actin-binding proteins (*Gas2l3, Flna, Coro6, Diaph a*nd *Ermn)*, a myosin (*Myh11*) and members of dynein family (*Dnah5, Dync1i1, Dynll1, Dnaic1, Dnah8)* (Fig. 4d, Supplementary table S1).

## Discussion

Familial forms of ALS with mutations in the *FUS* gene are characterised by a relatively early onset and a fast progression of the disease (reviewed in ^4,12,18^). Even so, despite a constant production of a pathogenic variant of FUS, spinal motor neurons of FUS-ALS patients function normally for decades (average age of FUS-ALS is ∼43 years ^12^). This suggest that by activating certain compensatory mechanisms neurons are able to neutralise toxicity of pathogenic variants of FUS until their accumulation, particularly when it is coupled with mislocalisation, reaches a critical threshold or the cell defence machinery is compromised by an external or internal stress. Several mouse models of ALS caused by modified expression of FUS display the same pattern of the disease progression: animals are presymptomatic for a long period of their postnatal life (3-12 months, depending of the introduced genetic modification) but after the manifestation of first clinical signs of pathology the disease progresses rapidly, mice develop severe motor dysfunction and die within several days (reviewed in ^10-12^). However, this phenotype is typical for mouse lines expressing high levels or/and highly pathogenic variants of human FUS. In several recently produced models a problem of artificially high levels of endogenous FUS expression was solved by using knock-in techniques ^40,41^, a relevant promoter ^38^ or selection of mouse lines with an appropriate level of transgene expression ^42^. Although the common trend for later onset and longer disease duration was observed, each model displayed distinct behavioural and histopathological characteristics.

Expression of human FUS carrying point mutations in the NLS (either FUS^R521C^ or FUS^R521H^) at the level comparable with the level of expression of the endogenous mouse FUS caused progressive motor dysfunction caused by the loss of muscle innervation, spinal motor neurons and their axons in the absence of FUS mislocalisation to the cytoplasm and formation of inclusions ^38^. In the spinal cord of ageing symptomatic transgenic mice in the absence of the endogenous mouse FUS the most prominent changes revealed by RNA sequencing analysis were downregulation of mRNA encoding proteins involved in protein synthesis and synaptic function, and upregulation of mRNA encoding chaperons and proteins involved in RNA metabolism although no changes in a global splicing pattern was observed.

In another recent study, human FUS lacking NLS was expressed under control of Thy-1 promoter in the nervous system of transgenic mice at the level of around 80% of the endogenous mouse FUS expression. These mice developed a progressive motor impairment from the age of 12 weeks and significantly decreased lifespan. Cytoplasmic accumulation of deltaNLS FUS and formation of large inclusions were observed in upper motor neurons before the onset of clinical signs but in the spinal cord motor neurons, only at the age of one year. Neuroinflammation and the loss of upper but not lower motor neurons was prominent in 1-year old mice. At the same age cerebral cortex transcriptome analysis identified a large number of genes up- and downregulated as the consequence of deltaNLS FUS expression but again, no notable changes of splicing pattern were found.

To reveal changes in gene expression taking place in the nervous system of mice carrying ALS-related modifications of FUS before they develop any clinical signs of pathology, Humphrey et al. (2019) compared results of transcriptomic analysis of neural tissues from homozygous late-gestation embryo (E17.5 spinal cord or E18.5 brain) of two independent FUS knock-in mouse lines expressing FUS lacking nuclear localisation signal ^40,41,43^. These data were compared to data obtained in the same studies by RNA sequencing analysis of similar neural samples of FUS knockout mice, which allowed to discriminate changes caused by the loss of FUS function and the gain of function by FUS lacking NLS and therefore partially mislocalised to the cytoplasm. A substantial overlap of DEGs in knockout and knock-in models suggested that at the presymptomatic stage the effects of the substitution of endogenous FUS to deltaNLS FUS on gene expression is due to the loss of function, i.e. an inadequate functionality of deltaNLS FUS that remains nuclear in neurons of studied mice. Between identified DEGs, the most prominent changes common for knockout and knock-in, as well as specific for homozygous knock-in mice were upregulation for those that code for proteins involved in RNA metabolism and downregulation of those that code for proteins involved in synaptic transmission. This is consistent with considerable changes of splicing pattern identified in these studies and the growing body of evidence that defects of synaptic transmission, particularly in the neuromuscular junctions, are very early events in the development of pathology in animal models and FUS-ALS.

In our previously described model, S-FUS[1-359] transgenic mouse line, expression of highly aggregation prone and pathogenic tr-hFUS protein caused a relatively early onset (at the age of ∼13-18 weeks), severe and fast progressing motor pathology despite its level in neural tissues at presymptomatic stage was lower than the level of endogenous FUS ^13^. In the spinal cord of mice of a new L-FUS[1-359] transgenic line described here the level of Thy-1 promoter-driven expression of the same tr-hFUS protein was even lower than in the spinal cord of S-FUS[1-359] transgenic line. Consequently, these animals did not develop motor deficiency, had the same lifespan as their wild type littermates and were phenotypically indistinguishable from them. We suggested that when the abundance of this pathogenic protein is low, neurons are able to neutralise any of its damaging effects by adjusting certain intracellular mechanisms and pathways. In an attempt to identify these adjustments, we analysed changes in the spinal cord transcriptomes of young adult L-FUS[1-359]. Importantly, in this model no pathology-driven changes in gene expression take place and therefore, any observed expression changes should be linked to adaptation to production of tr-hFUS.

Substantial changes in the gene expression profile (272 of DEGs, FDR < 0.05) were found in the spinal cord of L-FUS[1-359] mice although for most DEGs fold changes were relatively subtle (average log2 fold change for upregulated DEGs was 0.8 and for downregulated DEGs – 0.6). The majority of identified DEGs represented neuron-specific genes or genes expressed in neurons and other types of cells, suggesting that gene expression changes is direct effect of tr-hFUS expression in neurons. Minimal changes in expression of glia-specific genes is consistent with the absence of neurodegenerative process and reactive gliosis in the spinal cord of L-FUS[1-359] mice. Due to the deletion of RGG, Zn-finger and a part of RRM motifs, tr-hFUS lacks the ability to bind target RNAs and therefore cannot directly affect RNA metabolism. Therefore, it was not surprising that mRNA encoding proteins involved in RNA metabolism were barely represented in L-FUS[1-359] DEGs in contrast to high representation of these mRNA in DEGs identified in studies of mouse lines expressing low levels of FUS variants with intact RNA-binding domains ^38,40^ Humphrey et al., 2019).

Between gene clusters identified in our analysis of RNA sequencing data, the most prominent were upregulation of expression of genes encoding proteins involved in cell adhesion and structural organisation of extracellular matrix. These proteins function as regulators of cell migration, axon guidance and structuring of synaptic contacts, and therefore play important roles in the assembly and function of neuronal networks. Despite being predominantly nuclear protein, FUS has important functions in axonal and synaptic processes ^44-49^. It is feasible that the presence of tr-hFUS in these neuronal compartments might have a dominant negative effect on these function of the endogenous FUS. Increased production of proteins that can provide additional stabilisation of neuronal networks might be a compensatory mechanism that efficiently prevents potentially damaging effects of tr-hFUS. Consistently, we also found modified expression of several proteins involved in organisation of actin cytoskeleton and motor proteins, indicating that the remodeling of cytoskeletal structures and cellular transport take place in the spinal cord cell of L-FUS[1-359] mice.

Unexpectedly, expression of tr-hFUS protein did not prompt upregulation of genes encoding proteins involved in protein folding. Moreover, several chaperone or chaperonine encoding genes appeared to be downregulated in the spinal cord cell of L-FUS[1-359] mice, although only slightly. Probably at this level of expression tr-hFUS does not reach a concentration threshold required for its efficient aggregation and therefore, activation of protein folding mechanisms is not required.

Further detailed studies are required to explain prominent changes in expression of several genes encoding proteins involved in circadian rhythms because of different directions of these changes and complex feedback loops regulating expression of these genes.

Overall, we found that the pattern of gene expression changes in the spinal cord of asymptomatic mice expressing a low level of cytoplasmically mislocalised, aggregation prone and deficient in RNA binding variant of human FUS is substantially different from patterns previously reported for either symptomatic or presymptomatic mice expressing various human FUS variants at similarly low levels. Observed changes reflects efficient adaptation of spinal motor neurons to expression of a potentially pathogenic protein and therefore, molecular mechanisms and pathways affected by these changes might be considered as valid targets for preventive therapy of FUS-ALS.

## Materials and Methods

### Animals

The L-FUS[1-359] mouse line was produced by pronuclear microinjection of a transgenic cassette for expression of human FUS under control of regulatory elements of the mouse Thy-1 gene as described in our previous publications ^13,19^. The cassette was identical to the cassette used for production of S-FUS[1-359] mouse line and therefore, the same mouse genotyping protocol was employed ^13^. Mice were maintained under 12/12 hour dark/light cycle with free access to food and water. All animal work was carried out in accordance with the Rules of Good Laboratory Practice in Russian Federation (2016). The Bioethics committee of Institute of Physiologically Active Compounds, Russian Academy of Sciences provided full approval for this research (Approval No. 20 dated 23.06.2017).

### Comparison of transgene copy number by quantitative PCR

Genomic DNA from ear biopsies was purified with Wizard® SV Genomic DNA Purification System (Promega) and 0.1 µg was used for real-time qPCR amplification with primers 5’ – TCTTTGTGCAAGGCCTGGGT – 3’ and 5’ – TAATCATGGGCTGTCCCGTT – 3’ targeting human FUS in the transgenic cassette. Amplification of the GAPDH gene fragment with primers 5’ – CACTGAGCATCTCCCTCACA – 3’ and 5’ – GTGGGTGCAGCGAACTTTAT – 3’ was used as a reference when the relative number of transgenic cassettes in DNA samples was assessed. The CFX96 real-time PCR detection system (Biorad) and CYBR GreqPCR protocol were employed with the cycle parameters: 10 min at 95°C followed by 40 cycles of 15 s at 95°C and 60 s at 60°C. *Motor*

### behaviour testing

Animal motor performance was assessed on the inverted grid and accelerating rotarod as described previously ^20,21^.

A locomotor activity was monitored in a square 30 × 30 cm activity camera under 25 lux illumination using the TRU SCAN Activity Monitoring System (Coulbourn Instruments, Whitehall, PA, USA).

### RNA extraction

Total RNA was extracted from thoracic spinal cord of 9-week old male mice using RNeasy Plus Mini Kit with genomic DNA eliminator columns (Qiagen). For quantification and the quality control of RNA samples Qubit fluorimetry and TapeStation analysis were employed.

### Quantitative RT-PCR

1 μg of total RNA was reverse transcribed in the presence of random hexamer primers according to manufacturer’s instructions (Eurogene, Russia) and resulting cDNA was used for qPCR reaction with GAPDH as a reference gene as previously described ^22,23^. The same primers and amplification protocol as for the genomic qPCR protocol above were used.

### Protein extraction and Western blot analysis

Total proteins were extracted from the mouse spinal cords, separated by SDS-PAGE and transferred to PVDF membrane by semidry blotting as described elsewhere ^13,19^. Human FUS protein was detected using rabbit polyclonal antibody 14080 (a kind gift from Don Cleveland) specific to its N-terminal epitope, secondary anti-rabbit HRP-conjugated antibodies (GL Healthcare) and WesternBright TM Sirius chemiluminescent detection system (Advansta). For loading control membranes were re-probed with mouse monoclonal antibody against beta-actin (clone AC-15, Sigma-Aldrich).

### Histology and immunohistochemistry

The protocol for preparation of histological sections of the mouse spinal cord and immunodetection of human FUS with antibody 14080 was described previously ^13^.

### RNA sequencing

cDNA libraries for the dual indexed single-end sequence analysis were prepared from equal amounts (270 ng) of each total RNA sample using Illumina TruSeq Stranded Total RNA LT Sample Prep Kit. Following libraries’ quality checks and normalisation, cDNA libraries were sequenced on Illumina NextSeq 500 to generate single end 75 bp reads.

### Sequencing data analysis

Processing of RNA sequencing raw data included the adapter, length and quality trimming by Trimmomatic, mapping of reads to the mouse genome (release GRCm38) by STAR aligner, counting the overlap of reads with genes by featureCounts, implemented in PPLine script ^24-27^. Differential gene expression analysis was performed with the edgeR package using a Fisher exact test between experimental groups ^28^. We used the Benjamini-Hochberg method for multiple testing FDR correction. The genes with expression level ≥ 1 Log10 CPM (counts per million) and FDR ≤ 0.05 were taken into the account and considered as differentially expressed. Multidimensional scaling between all experimental samples were performed with limma package, pairwise distances were calculated using root-mean-square of Log2FoldChange values between experimental groups^29^. Sorting of microglial and neuronal genes of the whole spinal cord samples were performed using the published datasets of purified microglia and laser-microdissected ventral horns of mouse spinal cord ^30,31^. We applied a 5-fold change cutoff of gene expression levels to discriminate neurons- and microglia-specific genes as suggested previously ^31,32^.

The analysis of enriched gene ontology (GO) terms was performed using DAVIDWebService package for R with a P-value = 0.05 (Fisher exact test) and Q-value = 0.05 (BH-corrected p-value) as cutoffs ^33^. Only those GO terms that have enrichment by differentially expressed genes ≥ 4 were taken for further analysis. Then, redundant GO terms were removed using REVIGO software ^34^.

Differential expression analysis, data visualization, and GSEA (Gene Set Enrichment Analysis) were performed using R project for statistical computing ^35^. Visualization of experimental data was made with ggplot2 and GOplot R packages ^36,37^.

RNA sequencing data were deposited in Gene Expression Omnibus (GEO) under the number GSE130604.

## Acknowledgements

We are thankful to Angela Marchbank and Georgina Smethurst from the Cardiff School of Biosciences Genomics Research Hub for help with RNA sequencing, and Don Cleveland for sharing with us human FUS specific antibody. This study was supported by Russian Science Foundation projects RScF#18–15-00357 (phenotyping of L-FUS mice), RScF#17-75-20-249, (producing the L-FUS mice) and the Motor Neuron Disease Association research grant (Buchman/Apr13/6096). Bioresource Collection of IPAC RAS (No. 0090-2017-0016) and Core Facility IGB RAS were used to maintain transgenic mice and test their behaviour using equipment of the Centre for Collective Use IPAC RAS. RNA sequencing analysis was supported by Russian President Foundation grant MК-3316.2019.4. Differential expression analysis was performed using the equipment of the Engelhardt Institute of Molecular Biology RAS “Genome” Center (http://www.eimb.ru/rus/ckp/ccu_genome_c.php).

## Author Contributions Statement

V.L.B., S.O.B. and N.N. designed the study and wrote the main manuscript text; E.A.L. produced molecular biology data and prepared figures 2, 3 and S1; S.F. and A.P.R. provided sequencing data analysis and prepared figure 4 and supplementary table S1; K.D.C. analyzed protein by Western blot and prepared figure 1; M.S.K. analyzed ectopic hFUS by immunohistochemistry; A.A.U. and A.V.D. produced transgenic mice and prepared Supplementary figure S2; I.A.F. and S.B. defined chromosomal localization of transgenic cossets in both hFUS[1-359] lines.

## Additional Information Statement

The authors declare no competing interests.

