## Supplemental Figures for "Low level of expression of C-terminally truncated human FUS causes extensive changes in spinal cord transcriptome of asymptomatic transgenic mice"

**Supplementary Information**

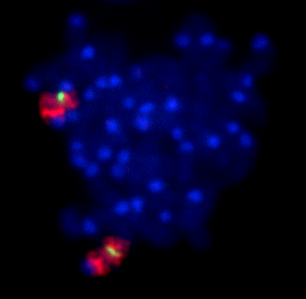

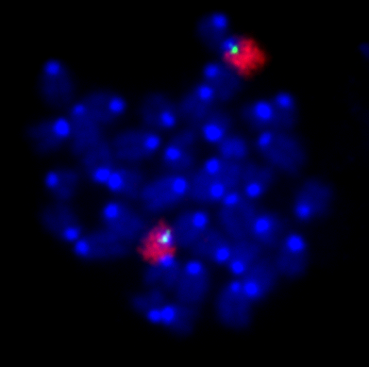

**S-FUS[1-359]**

**L-FUS[1-359]**

**mChr12-specific paint probe**

**mChr11-specific paint probe**

**transgenic cassette probe**

**transgenic cassette probe**

**b**

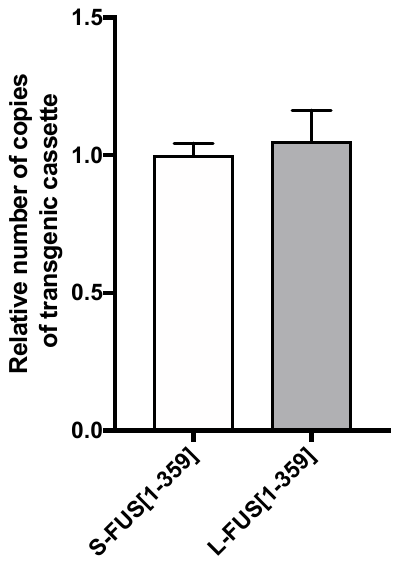

**a**

**Supplementary Figure S1.** a) Relative number of copies of transgenic cassettes integrated in the

genome of hemizygous L-FUS[1-359] and S-FUS[1-359] mice assessed by qPCR analysis of genomic DNA extracted from ear biopsies. b) Tandemly arranged transgenic cassettes mapped by FISH to chromosome 11 in L-FUS[1-359] and chromosome 12 in S-FUS[1-359] mice. In situ hybridisation of homozygous mouse nucleated blood cell metaphase plates with transgenic cassette DNA labelled with green496-dUTP (Enzo life sciences) by nick translation and biotin-labelled mouse chromosome specific paint probes (Cambio) that were detected using Texas red conjugated antibodies.

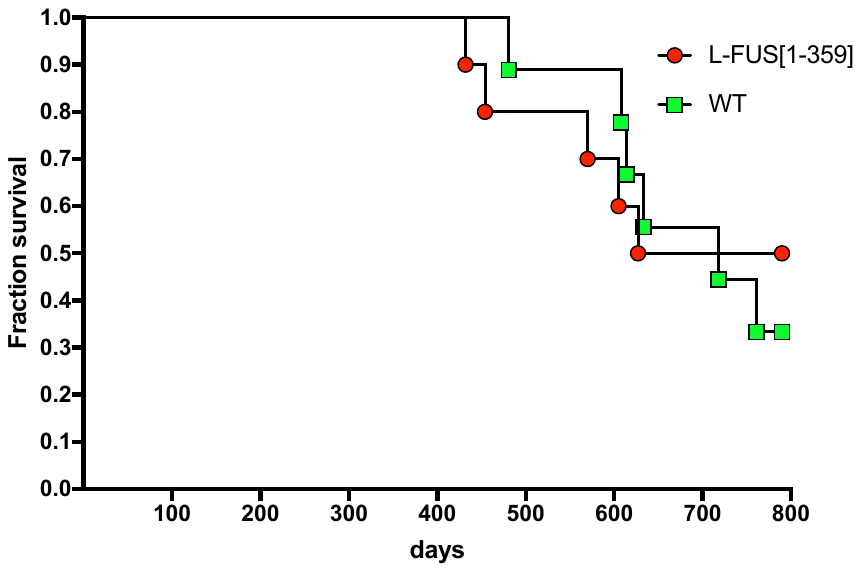

**Supplementary Figure S2.** Kaplan-Meier survival plot for wild type (WT) and homozygous male L-FUS[1-359] mice. The experiment was censored for 26 months. No statistically significant difference was found between two groups by log rank test.

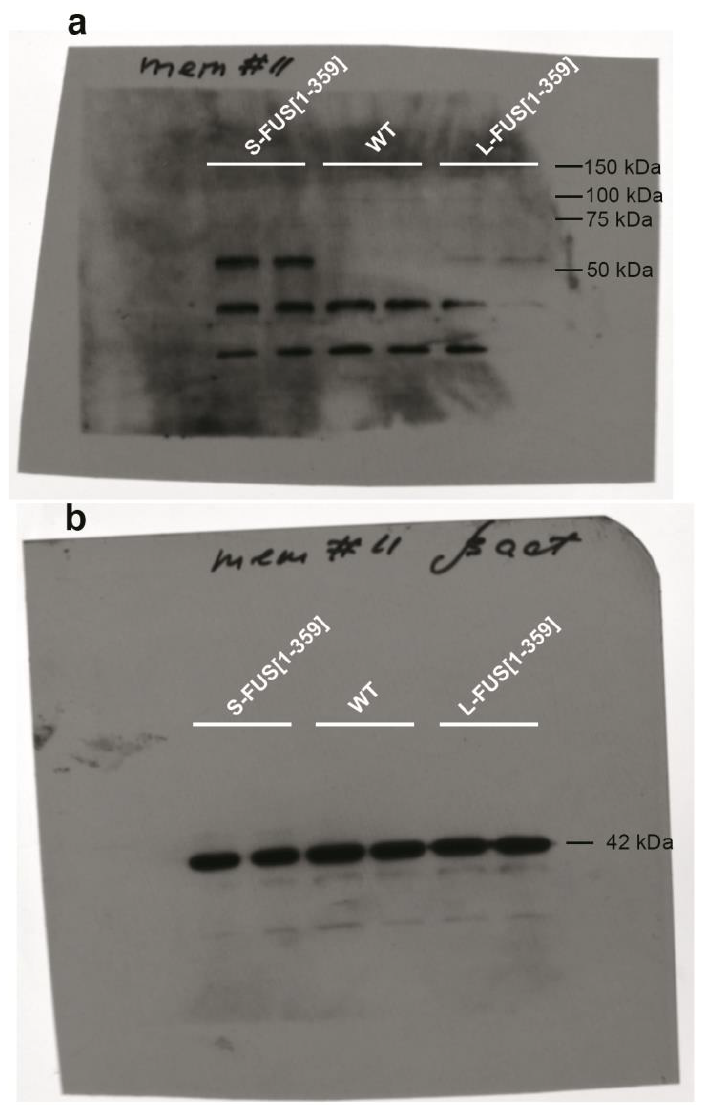

**Supplementary Figure S3.** Full-length images of Western blots used for figure 1b. Total proteins have been separated by 10% SDS-PAGE, transferred to PVDF membrane and probed with (a) rabbit polyclonal antibody 14080 specific to N-terminal epitope of human FUS protein. Secondary anti-rabbit HRP-conjugated antibodies (GL Healthcare) were used followed by chemiluminescent detection with WesternBright TM Sirius (Advansta) and 30 seconds exposure with X-ray film. The same membrane was re-probed with (b) mouse monoclonal antibody against beta-actin (clone AC-15, Sigma-Aldrich) and secondary anti-mouse HRP-conjugated antibodies (GL Healthcare).
